# A Simultaneous EEG-fMRI Study of Thalamic Load-Dependent Working Memory Delay Period Activity

**DOI:** 10.1101/2020.10.16.342568

**Authors:** Bernard A. Gomes, Chelsea Reichert Plaska, Jefferson Ortega, Timothy M. Ellmore

## Abstract

Working memory (WM) is an essential component of executive functions which depend on maintaining task-related information online for brief periods in both the presence and absence of interfering stimuli. Active maintenance occurs during the WM delay period, the time between stimulus encoding and subsequent retrieval. Previous studies have extensively documented prefrontal (PFC) and posterior parietal (PPC) cortex activity during the WM delay period, but the role of subcortical structures including the thalamus remains to be fully elucidated, especially in humans. Using simultaneous EEG-fMRI, we investigated the role of the thalamus during the WM delay period following low and high memory load encoding. During the delay, participants passively viewed scrambled images containing similar color and spatial frequency to serve as a perceptual baseline. Using individual fMRI-weighted source analyses centered around delay period onset, the effects of increased and decreased memory load on maintenance were observed bilaterally in thalamus with higher source activity evoked during low compared to high load maintenance. The main finding that thalamic activation was attenuated during high compared to low load maintenance suggesting a sensory filtering role for thalamus during consolidation of stimuli in WM where the highest evoked activity occurs when fewer stimuli need to be maintained in the presence of interfering perceptual stimuli during the delay. The results support the idea that the thalamus plays a role in short-term memory maintenance by regulating processing of interfering stimuli.

## Introduction

Models of memory differentiate between sensory memory, measured in milliseconds to seconds; short-term memory or working memory, which persist from seconds to minutes; and long-term memory, which may last for decades (Atkinson, Brelsford, & Shiffrin, 1967). Working memory (WM) is the ability to maintain and manipulate information for the guidance of goal-directed behavior (A Baddeley & Hitch, 1974). WM encompasses storage and processing functions like active maintenance of goal-related information, thinking, and planning (Alan Baddeley, 2003). Early work by Fuster & Alexander (1971) and others in nonhuman primates have revealed that neurons in the PFC show elevated levels of action potential firing during the maintenance phase of delayed-response tasks (Funahashi, Bruce, & Goldman-Rakic, 1989). The maintenance phase is called the delay period and is characterized by elevated neural activity (also referred to as delay activity) that occurs during maintenance of information, or the period between encoding and retrieving a stimulus. This neural signature represents the temporary memory storage of the stimulus in WM (Fuster & Alexander, 1971). Current research focuses on activity during the WM delay period after visual stimuli are encoded, but before memory recall.

Fuster & Alexander (1971) first reported that changes were observed during the delay period in a short-term memory task in the prefrontal cortex (PFC) and mediodorsal nucleus of the thalamus (MDt). Although the longstanding view of the thalamus is that it serves as a relay station for all major sensory pathways, it has more recently been suggested that the thalamus plays a role in memory and cognition by maintaining and updating relevant information (Wolff & Vann, 2019). Electrophysiological recordings in animal studies have suggested that MDt is strongly involved in WM (Mair et al., 2015; Sommer & Wurtz, 2006). MDt receives inputs from parahippocampal regions and is also reciprocally connected to the medial prefrontal cortex (mPFC) (Jang & Yeo, 2014; Yang, Logothetis, & Eschenko, 2019). These connections between the thalamus and PFC suggest the importance of thalamus in WM by helping activity persist in PFC through the connections via MDt (McCormick & Bal, 1994; Yang et al., 2019). Previous research has suggested that the neural activity generated during the delay period is maintained in the cortex, particularly the anterior lateral motor (ALM) cortex (Courtney, Ungerleider, Keil, & Haxby, 1997; Inagaki, Fontolan, Romani, & Svoboda, 2019). However, the idea has been challenged recently, where a study found that the maintenance of information is dependent on delay activity in the thalamus (Guo et al., 2017). Specifically, the MDt sends connections to the PFC, a region that is implicated in WM maintenance (Mitchell, 2015). More recently, MDt and the anterior thalamus has been shown to play a role in familiarity and recollection, respectively (Kafkas, Mayes, & Montaldi, 2019). In the same study, it was also reported that ventral posteromedial and pulvinar thalamic nuclei regions were involved in scene familiarity such that there was greater activity for familiar scenes when compared to new scenes. These findings suggest that thalamus is implicated in WM.

Recent neuroimaging studies have also reported differential activation in Default Mode Network (DMN) regions during various cognitive tasks, including WM (Koshino, Minamoto, Yaoi, Osaka, & Osaka, 2014). It is important to consider the role of DMN in WM, which has been implicated in memory and attention tasks such that DMN regions show decreased activity during demanding cognitive tasks (Shulman, Corbetta, Buckner, Fiez, et al., 1997), and is also linked to the thalamus (Wang et al., 2014). The DMN consists of distinct regions that are more active at rest than during the task. The regions include the dorsolateral prefrontal cortex, inferior frontal cortex, left inferior temporal gyrus, retrosplenial cortex, medial frontal regions, amygdala, and the more recently added subcortical structures like thalamus and basal forebrain (Mazoyer et al., 2001; Shulman, Corbetta, Buckner, Raichle, et al., 1997). The DMN is known to be involved in mind-wandering and lack of awareness of external space (Christoff, Gordon, Smallwood, Smith, & Schooler, 2009). DMN has also been implicated strongly in cortical integration that allows transmodal information processing that is not related to the immediate sensory input (Alves et al., 2019; Margulies et al., 2016). The DMN is typically active at rest, that is when a participant is not engaged in a task, but it has also been implicated in non-rest cognitive processes like WM (Pyka et al., 2009). Recent investigations also highlighted the DMN’s role in WM and episodic memory tasks suggesting differential activation during different memory phases (Daselaar et al., 2009; Woodward, Feredoes, Metzak, Takane, & Manoach, 2013). Therefore, we were interested in testing whether there is a differential load-dependent activation pattern in the cortical delay activity with thalamus as the source. We expected to find an increase in thalamic activity as load increases because it helps neural activity persist during the delay activity. Conversely, if the thalamus, as a part of DMN, modulates WM during delay activity, we expected less thalamic activation as load increases.

The anatomical components of the memory system include the hippocampus and various structures interconnected with the hippocampus. The structures include the surrounding entorhinal cortex, perirhinal cortex, parahippocampal cortex, amygdala, mammillary bodies, and anterior thalamic nuclei. Parahippocampal (PHC) regions have been implicated in the encoding processes of memory, particularly scene memory (Alkire, Haier, Fallon, & Cahill, 1998; R. Epstein, Graham, & Downing, 2003; Maguire, Frith, Burgess, Donnett, & O’keefe, 1998).

In an event-related functional magnetic resonance imaging (fMRI) investigation, researchers found that both the left and right parahippocampal gyrus strongly respond during encoding and retrieval (Rombouts, Barkhof, Witter, Machielsen, & Scheltens, 2001). Furthermore, a recent fMRI investigation also reported a memory load effect observed in the PHC during WM encoding of spatial layouts (Schon, Newmark, Ross, & Stern, 2015). In this research study, we were also interested in testing whether there is a differential load-dependent activation pattern in the parahippocampal gyrus during encoding. PHC’s role as a WM buffer for scene processing and mnemonic encoding of novel scenes is well-established (R. Epstein, Harris, Stanley, & Kanwisher, 1999; Preston et al., 2010). Accompanying the view of PHC being a WM buffer for scene memory (R. A. Epstein, Parker, & Feiler, 2007), we expected the region to show sustained activity during encoding, which is greater with the higher memory load. Results from behavioral studies suggest that stimuli are subjected to transient encoding after presentation to be actively maintained in the absence of bottom-up simulation (Jolicœur & Dell’Acqua, 1998; Raye, Johnson, Mitchell, Reeder, & Greene, 2002). Furthermore, an event-related fMRI study found that parahippocampal areas are recruited during WM encoding of scenes (Ranganath, DeGutis, & D’Esposito, 2004). Therefore, an additional aim of the study is to test whether there will be differential activation in the PHC regions during encoding as a function of memory load using source analysis.

In this study, we collected simultaneous electroencephalography (EEG) and functional magnetic resonance imaging (fMRI) from 24 subjects. Due to the complementary strengths of each of the recording modalities, combining EEG-fMRI harnesses both high temporal and spatial resolution. The goal of the study was to examine the spatiotemporal patterns that underlie the encoding and maintenance of visual memory using a modified Sternberg scene working memory task (Sternberg, 1966). The task involved two memory loads, a low-load with two scenes presented sequentially during encoding, and a high-load with five scenes presented, in order to study the difference in cognitive demand during maintenance of complex naturalistic visual information. We computed event-related potentials (ERP) during the delay and encoding periods to examine the differences between the task conditions (low-vs. high-load) and used source analysis weighted by the individual’s fMRI activity to constrain the potential sources of these signals.

We hypothesized that the thalamus would be sensitive to the difference in load or the amount of information maintained during the delay period because of its theorized role in helping neural activity persist in connected cortical regions like PFC. Memory load has been extensively studied to understand if there are limits in how much information can be maintained in WM and how the brain can maintain multiple items in WM (Fukuda, Awh, & Vogel, 2010; Sweller, 2011). These studies have found that during the delay period, the dorsolateral prefrontal cortex (dlPFC) and the middle and superior frontal gyri show increased activity as memory load increases. In contrast, left caudal inferior frontal gyrus shows increased activity as memory load decreases (Manoach et al., 1997; Rypma, Berger, & D’esposito, 2002; Rypma, Prabhakaran, Desmond, Glover, & Gabrieli, 1999). Thalamic relays to and from dlPFC have been suggested to suppress interfering environmental stimuli during delay activity (Postle, 2005). During the delay period, the thalamus may be regulating sensory processing by up-or down-regulating potentially disruptive sensory information (Knight, Staines, Swick, & Chao, 1999). The disruptive sensory information in our study was served by presenting scrambled images during delay, which contained similar color and spatial frequency as the scene stimuli presented during the encoding period.

## Materials and Methods

### Participants

A total of 24 participants were recruited between August 2017 and August 2018 by flyers posted throughout City College of New York campus. The study consisted of healthy adults between ages 18 and 54 with normal or corrected-to-normal vision and the ability to make button presses. Participants were excluded if they had a history of neuropsychological disorders. Each participant provided written consent and completed the study procedures according to a protocol approved by the City University of New York Institutional Review Board. Participants were either compensated $15 per hour of participation or one extra course credit per hour of participation.

All the participants were included in the fMRI analysis (12 males, 12 females, age range 18 to 54, mean age 25.3 years, SD = 8.5). For the EEG analysis, a total of 2 participants were excluded. One participant had excessive noise in their EEG signals and another participant failed to remain awake during the task. The final sample included in the EEG analyses was 22 subjects.

### Task Design and Stimuli

EEG and fMRI data were acquired simultaneously in a single experimental session. The participant completed a modified version of a Sternberg WM task (Sternberg, 1966). The task consisted of an encoding phase (2 or 5 scene images presented for 1400 msecs each), delay phase (6 scrambled images, 6000 msecs), and a probe recognition phase (1400 msecs), followed by an average 3000 msecs jitter period (*Figure 1*).

**Figure 1:**
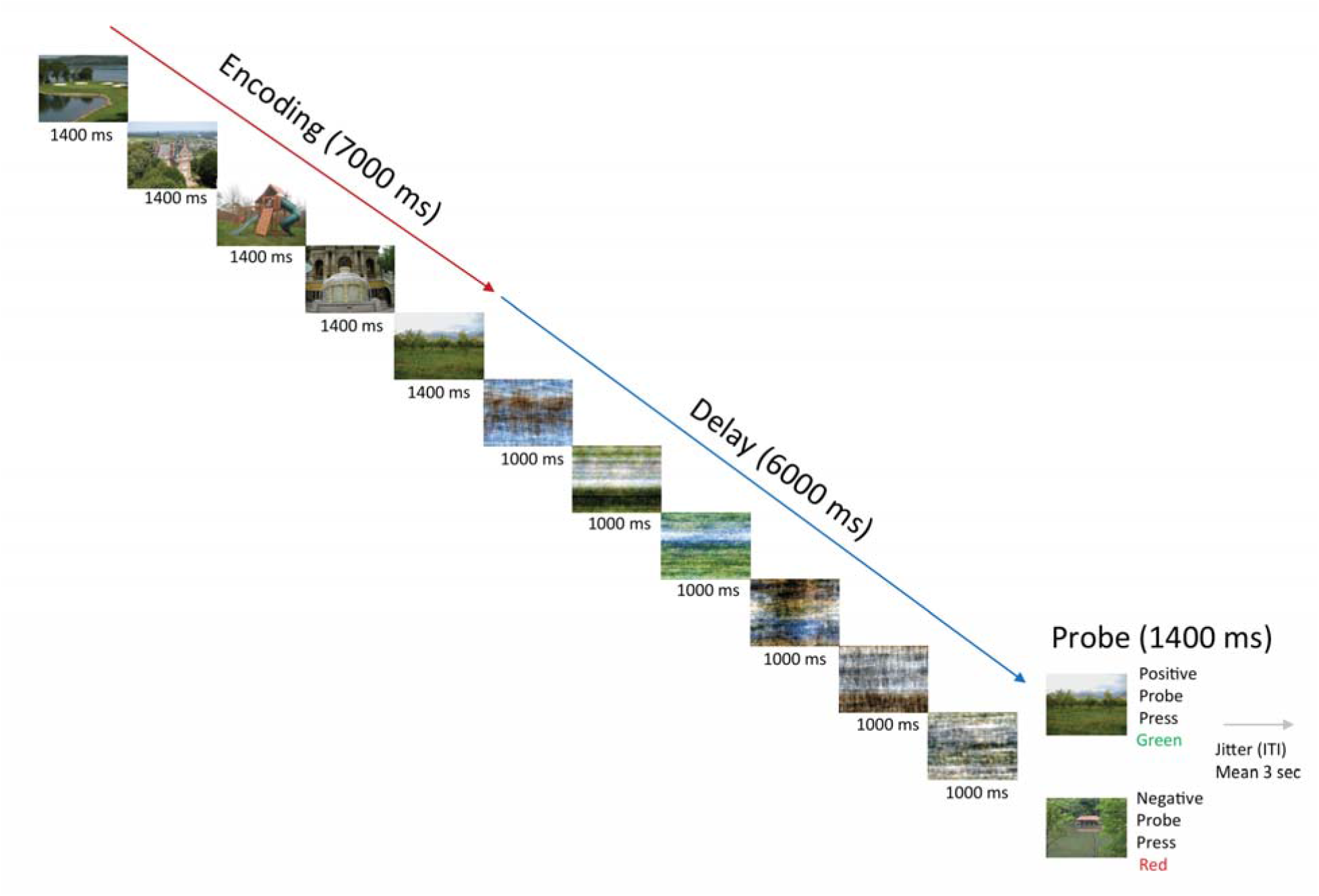
Task Design for the Working Memory Task. Subjects performed a modified Sternberg task with naturalistic scenes consisting of 50 trials per working memory load. There were two working memory loads: low load-2 images and high load-5 images. An example of high-load trial is shown below. Subjects viewed either 2 or 5 sequentially presented images (encoding phase), maintained the scenes across a 6-s length delay period (maintenance phase), and determined whether a probe scene matched one of the previous images seen during that trial or was a nonmatch (test phase). Each trial ended with a variable-length fixation/ITI (Mean = 3 sec).

During the task, each participant completed 50 trials of low load (2 scene images) and 50 trials of high load (5 images) trials presented in separate runs with order randomized and counterbalanced. In each run, the encoding phase was followed by the delay phase, where the participant viewed Fourier phase-scrambled stimuli with similar color and spatial frequency. The phase-scrambled scenes provided a visual perceptual baseline while the participants maintained the scenes presented during encoding. Each trial ended with a recognition probe phase, with the participant making a choice on whether they had previously seen the image during the encoding period or not by making a button response. The scenes were randomly selected from the SUN database (Xiao, Hays, Ehinger, Oliva, & Torralba, 2010) consisting of 671 novel color outdoor scenes. Images were presented in 800 by 600 pixel format on an LCD monitor. A Magnetic Resonance Imaging-compatible amplifier (BrainAmp-MR, Brain Products, Germany) was used for the simultaneous EEG-fMRI recordings, which were taken directly inside the MRI bore with stimuli presented directly behind the subject’s head and viewable by a mirror attached to the head coil. Digitized button press signals were sent via fiber optic cable to the USB interface located in the control room. This set up ensured that there were no artifacts accumulated along the way to acquisition computers in the scanner control room. The short lengths of the electrical cables used to connect the electrode cap with the amplifier fulfilled all subject safety requirements without compromising data quality.

Each session consisted of an initial practice run and three experimental runs. The practice session took place outside the scanner and consisted of 3 trials per condition. The experimenter read task instructions from a script for all participants before the participant entered the scanner and before every run while inside the scanner. We did not instruct participants when to blink during the task. Blink artifacts were confirmed by visual inspection later during processing and removed from EEG data before averaging and source analysis.

### EEG Data Acquisition and Pre-processing

EEG was recorded with BrainAmp-MR, BrainProducts, Germany placed inside the MR scanner and sampled at 2500 Hz. Subjects were fitted with a MR-compatible EEG cap (BrainCap-MR 32 Channel-Standard, BrainProducts, Germany) containing a 10-10 montage with 32 electrodes including 31-scalp electrodes (Fp1, Fp2, F7, F3, Fz, F4, F8, FC5, FC1, FC2, FC6, T7, C3, Cz, C4, T8, CP5, CP1, CP2, CP6, TP9, TP10, P7, P3, Pz, P4, P8, POz, O1, Oz, O2) and one electrode for ballistocardiogram (BCG), placed on the left shoulder-blade. The impedances were kept below 20 kOhm and were monitored to stay below 50 kOhm during recording for safety. All the electrodes were referenced to one site, Fpz, during data collection, and all the electrodes on the scalp were re-referenced to the common average reference offline. The EEG was analyzed with Brain Electrical Source Analysis (BESA) Research v7.0. MRI artifacts were removed in BESA Research (Allen, Josephs, & Turner, 2000). The parameters used for fMRI artifact removal were 16 artifact occurrence averages on fMRI data with Repetition Time (TR) of 2000 msecs. The correction was done using either the MR pulse trigger or phase synchronization between EEG equipment and MRI scanner (Allen et al., 2000). The data were low pass filtered at 70 Hz for the artifact cleaning. All the data were down sampled from 1000/2500 Hz to 500 Hz for comparison.

Eye-blink artifacts and BCG artifacts were removed using defined topographies for correction. A data block containing the artifact was marked and either defined as an eye-blink or BCG. Then the BESA pattern matching algorithm selected the ICA channel that matched the highest explained variance (~95%) and subsequently used PCA to remove the artifact (Berg & Scherg, 1994; Moosmann et al., 2009). For eye-blink correction, the data were filtered between 1 to 12 Hz. For BCG correction, the data were filtered between 1 and 20 Hz, and a zero-phase filter slope was used. For low cutoff, the filter type was set at 12 dB/oct and for high cutoff, the filter type was set at 24 dB/oct.

Data were visually inspected, and muscle artifacts were removed by trained research assistants. Exceptionally noisy channels were interpolated for the electrode channels that proved problematic during data collection. The baseline was defined using the 100 msecs preceding the onset of the stimuli for each trial. For generating ERPs, the low cut off filter of 0.1 Hz was applied. After generating the ERPs, the blink and BCG artifacts were removed and a high cut off filter of 40 Hz was applied. The mean number of images across all trials that went into averaging was 277 (SD = 30) and 282 (SD = 13) for low-and high-load delay conditions, respectively. For low-and high-load encoding conditions, the mean number of images across all trials that went into averaging was 92 (SD = 8.3) and 236 (SD = 15.5), respectively. A paired sample t-test showed non-significant differences between delay conditions (p=0.354) and significant differences between encoding conditions (p<0.001). The latter result is because more images went into the averaging of high-load conditions (5 images) than low-load conditions (2 images).

### ERP Analysis

The EEG signal was segmented in epochs around stimulus onset for 1000 msecs at the start of the encoding period for the encoding period analysis and at the start of the delay period for the delay period analysis. Then the artifact-corrected epochs were averaged for each condition (high and low load) and task type (encoding and delay). The average ERPs for each condition were then used as input for group statistical ERP analysis performed with BESA Statistics v2.0 with appropriate multiple corrections across space and time using random permutation testing (Maris, 2012).

### Magnetic Resonance Imaging

While EEG was simultaneously recorded, subjects participated in a single one hour and thirty minute magnetic resonance imaging (MRI) session during which ten different MRI acquisitions were collected in the following order: 1) a five minute eyes open rest scan (functional), 2) either a high (N=5) or low (N=2) load working memory task (functional) with the high or low load order randomized across subjects, 3) a five minute eyes open rest scan (functional), 4) a T1-weighted volume (structural), 5) a T2-weighted volume (structural), 6) a five minute eyes open rest scan (functional), 7) either a high (N=5) or low (N=2) load working memory task (functional) depending on which load was presented during the second acquisition, 8) a five minute eyes open rest scan (functional), 9) a recognition task (functional) for old scenes presented in the low and high WM tasks mixed with new scenes not previously viewed, and 10) a PETRA volume (structural) to aid in electrode visualization and localization. Functional data acquired during the eyes open rest scans and during the memory tasks consisted of blood oxygen level dependent echo planar images (BOLD-EPI) with echo time (TE) of 30 msecs, repetition time (TR) of 2000 msecs, a field of view (FOV) of 249 mm, 35 axial slices, with 3 mm isotropic voxels. The T1-weighted structural was collected with a TE of 2.12, a TR of 2400, a 254 mm FOV with 1 mm isotropic voxels. The T2-weighted structural was collected with a TE of 408, a TR of 2200, and a 254 mm FOV with 1 mm isotropic voxels. The PETRA structural was collected with a TE of 0.07, a TR of 3.61, a 298 mm field of view with 0.938 mm isotropic voxels.

### Image Analysis

MRI data were processed using AFNI (Analysis of Functional Neuro Images: afni.nimh.nih.gov) (Cox, 1996). Each BOLD-EPI 4-dimensional volume timeseries was aligned to the T1-weighted volume using AFNI’s *align_epi_anat.py* python script, which also performed skull-stripping of the T1 volume, EPI slice timing correction, alignment of the EPI to the T1 using a 12 parameter affine transformation, and spatial blurring of the EPI timeseries using a gaussian full width at half maximum of 4 mm. First (subject) level statistical analysis of the processed individual subject EPI timeseries was performed using AFNI’s 3dDeconvolve with encoding (scenes) and delay (scrambled) periods of the working memory task modeled using 7 sec for the high load encoding period (2 sec for low load encoding) and 6 sec for delay period blocks respectively to form regressors which were convolved using a hemodynamic response function (HRF) of the form HRF(t) = int(g(t-s), s=0..min(t,d)) where g(t) = t^q * exp(-t) /(q^q*exp(-q)) and q = 4 before being added as regressors of interest to the general linear model design matrix. General linear tests were used to compare encoding (viewing scenes) to delay (viewing scrambled while maintaining previously viewed scenes). Regressors of no interest in the design included p=7 polynomial functions to model baseline shifts with a cutoff of (p-2)/D Hz where D is the duration of the imaging run and the three translation and three rotational subject motion parameters. Second (group) level statistical maps were computed using AFNI’s 3dttest++ by inputting each subject’s voxelwise regression coefficient maps to compare encoding (scenes) vs. delay (scrambled) for each load. The first level general linear test comparisons of encoding (scenes) vs. delay (scrambled) for high and for low WM load were output as individual subject maps in Talairach space with 2 mm isotropic resolution and thresholded using a false discovery rate of q=0.01 before being imported into BESA Research 7.0 as weight maps for constrained dipole source analysis (Scherg, 1990).

### Dipole Source Analysis

For each participant, the positions of the 32-channel electrodes used for simultaneous EEG-fMRI scanning sessions were estimated initially using an approximation of electrode locations made from a standard montage template (BESA-MRI-Standard-Electrodes) and then adjusted manually based on visual inspection of the indentation-artifacts caused by electrode on the scalp, which appeared like dips on the scalp in the surface reconstructions. An example of electrode locations in a single subject is shown in *Figure 2*. Further, for each participant, the anatomical MRI was segmented manually in BESA MRI v2.0 to create 4-layer Finite Element Model (FEM) realistic head models to be used in the source analysis. On the basis of individual electrode coordinates and landmark segmentation in Talairach Space, BESA calculated the best fitting ellipsoid of each subject (Scherg, 1992). For individual source analysis, fMRI statistical maps were imported for each condition and participant, followed by the process of dipole modeling. Seed-based functional analysis has been previously shown to be better modeling for analyzing subcortical structures like the amygdala, striatum, and thalamus (Bzdok, Laird, Zilles, Fox, & Eickhoff, 2013). Seed-based dipole fitting was based on *a priori* hypothesis testing to explain ERP changes in all conditions. For encoding conditions, two equivalent dipoles were fitted onto the bilateral parahippocampal cortex (PHC) for each participant in the two conditions. For delay conditions, two equivalent dipoles were fitted onto bilateral thalamus. A time window from each participant’s data window was chosen from onset to the peak of the first Global Field Potential (GFP) peak, which is a measure for spatial standard deviation as a function of time (Strik & Lehmann, 1993). An example of a subject’s GFP waveform is shown in *Figure 3* and an example analysis window used in source analysis is shown in *Figure 4*.

**Figure 2:**
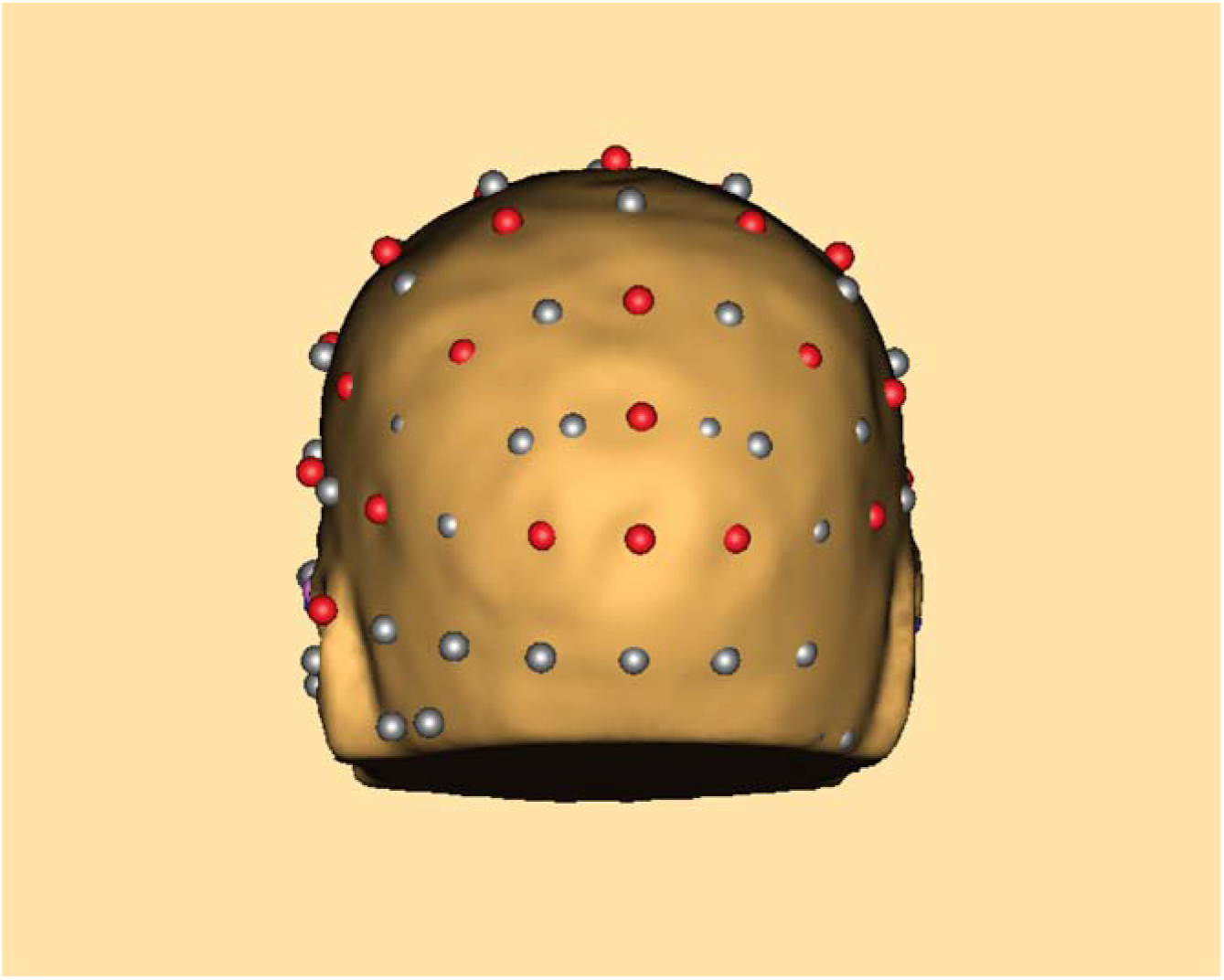
Example of electrode-localization superimposed on a single-subject scalp surface reconstruction. Standard set of 3D electrode positions based on the 10-10 electrode system was used to coregister electrodes with individual scalp surface reconstructions. Electrode positions were visually inspected to make sure the electrodes were as close to the indentation artifacts on the scalp caused by the electrode gel. An example from a single subject is shown above.

**Figure 3:**
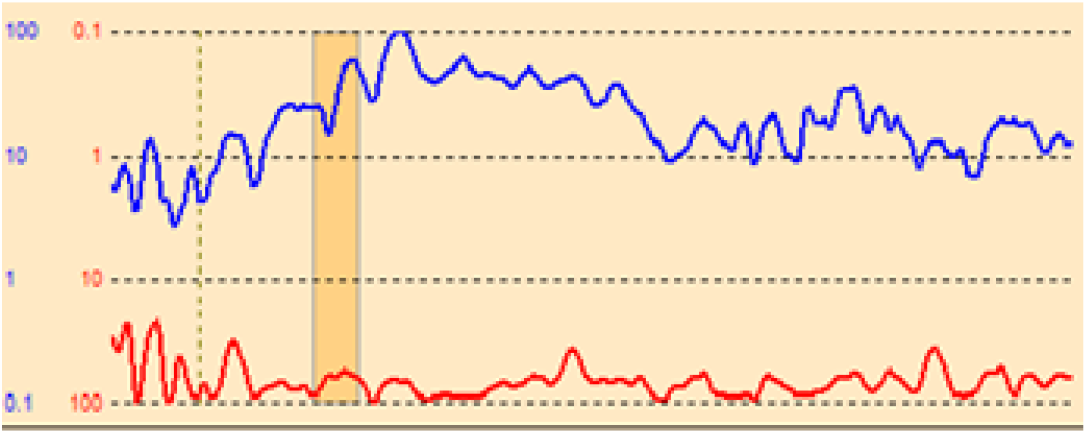
Example Global Field Power for Source Analysis. GFP (Global Field Power) of the original waveform of a single subject representing all combined delay activity of the experiment (blue) is displayed in logarithmic scale. The unexplained fraction of the data variance, or Residual Variance (RV) is also displayed (red) in inverted logarithmic scale. The fit process finds a source model that minimizes this RV. RV = 62% and length of duration of the x-axis is 1000 msecs.

**Figure 4:**
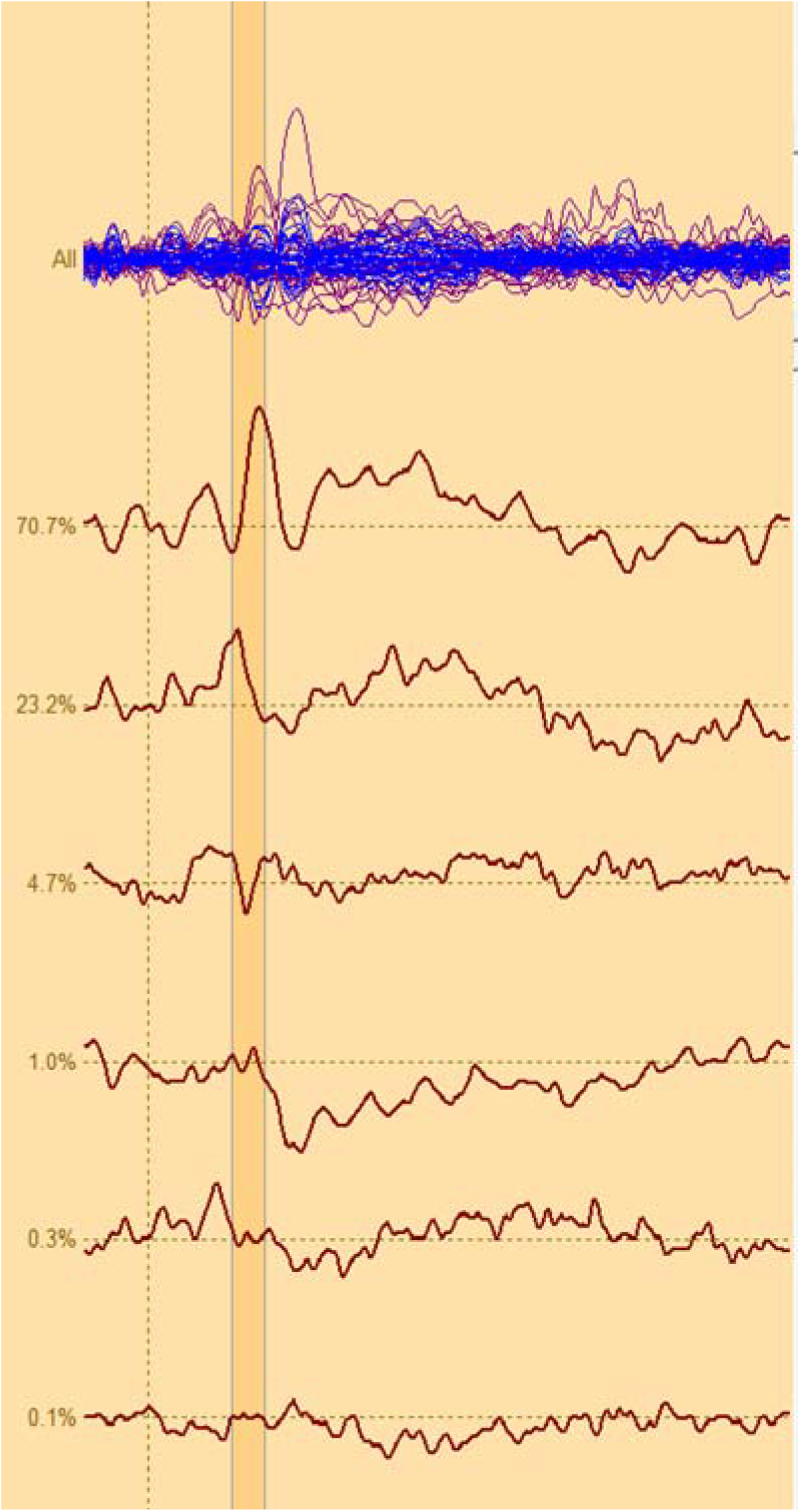
Example analysis window used in the source analysis. The upper trace shows the butterfly-plot of the averaged spike-signal (epoch duration: - 100 msec to 998 msec; the dotted line represents the stimulus onset). The model waveforms generated by the current source model is shown in blue. The following traces show the sourcewaveforms corresponding to the detected component (highlighted); numbers on the left indicate the contributed variance of each component to the measured signal. Note that one component accounts for more than 70% of the signal on the ascending slope.

During the seeding of dipole locations, fMRI activation maps were initially turned off to avoid potential bias in determining initial seed location. The dipoles were then fit onto the respective sources weighted by the fMRI statistical maps using the RAP-MUSIC algorithm as implemented in BESA source space that estimates the dipole locations using the weighted MRI images (Grech et al., 2008). The dipole positions were constrained to stay within the target regions, but their orientations were kept free before the fit. All the dipoles fell within the appropriate brain regions (PHC and thalamus) after the fit. The dipole positions were expressed as Talairach coordinates in units of millimeters (mm) and averaged across all subjects. For delay conditions, the Talairach coordinates for left hemisphere dipole were x=-13.4, y=-21.9, z=3.7, and right hemisphere dipole coordinates were x=11.6, y=-21.8, z=3.8. Both dipole coordinates during delay always fell within the thalamus for all participants. For encoding conditions, the Talairach coordinates for the left hemisphere were x=-25.3, y=-38.1, z=-9.6, and right hemisphere dipole were x=24.7, y=-37.9, z=-9.4. Both dipole coordinates during encoding always fell within the PHC for all participants. *Tables 1 and 2* lists the individual coordinates for all the participants included in the source analysis during delay and encoding conditions, respectively. The source waveforms for each participant and condition were exported and then imported for group source analysis in BESA Statistics v2.0.

**Table 1:** Individual Talairach Coordinates (mm) during the Delay Condition

|  | **Right** |  |  | **Left** |  |  |
| --- | --- | --- | --- | --- | --- | --- |
|  | **x** | **y** | **z** | **x** | **y** | **z** |
| Participant 1 | 13.4 | -25.3 | 0.5 | -18.8 | -25.6 | 1.6 |
| Participant 3 | 13.4 | -25.3 | 0.5 | -17.8 | -25.2 | 1.2 |
| Participant 5 | 13.4 | -25.3 | 0.5 | -15.6 | -27.4 | 0.5 |
| Participant 6 | 13.4 | -25.3 | 0.5 | -15.6 | -25.6 | 4 |
| Participant 7 | 13.4 | -25.3 | 0.5 | -18.8 | -25.3 | -0.5 |
| Participant 8 | -6 | -9 | 6 | -6 | -9 | 6 |
| Participant 9 | 9.1 | -14.5 | 1.6 | -15.6 | -14.5 | 0.5 |
| Participant 10 | 17.8 | -26.8 | 8.5 | -10.2 | -27.4 | 4.8 |
| Participant 11 | 8.1 | -15.3 | 4.8 | -11.3 | -15.3 | 2.7 |
| Participant 12 | 15.1 | -21.7 | 2.8 | -15.6 | -21.7 | 5.9 |
| Participant 13 | 14.5 | -26.4 | 5.9 | -13.4 | -26.4 | 4.8 |
| Participant 14 | 16.1 | -30.7 | 2.6 | -14.5 | -30.7 | 3.8 |
| Participant 15 | 12.4 | -21.2 | 4.8 | -14.3 | -21.2 | 3.8 |
| Participant 16 | 6 | -9 | 6 | -6 | -9 | 6 |
| Participant 17 | 12.5 | -29.5 | 2.9 | -15.3 | -29.8 | 4 |
| Participant 18 | 6 | -9 | 6 | -6 | -9 | 6 |
| Participant 19 | 12.4 | -21 | 4.8 | -13.4 | -21 | 4.8 |
| Participant 20 | 19.2 | -33.1 | 3.2 | -14.1 | -33.2 | 4.3 |
| Participant 21 | 14.5 | -27.4 | 4.8 | -18.8 | -27.4 | 3.8 |
| Participant 22 | 11.3 | -19.9 | 0.5 | -10.2 | -19.9 | 1.7 |
| Participant 23 | 9.1 | -17.8 | 4.8 | -9.1 | -17.8 | 5.9 |
| Participant 24 | 10.2 | -19.9 | 4.8 | -13.4 | -19.9 | 4.8 |
| **Average:** | **11.6** | **-21.8** | **3.5** | **-13.4** | **-21.9** | **3.7** |

**Table 2:** Individual Talairach Coordinates (mm) during the Encoding Condition

|  | **Right** |  |  | **Left** |  |  |
| --- | --- | --- | --- | --- | --- | --- |
|  | **x** | **y** | **z** | **x** | **y** | **z** |
| Participant 1 | 25.3 | -35 | -10.2 | -25.3 | -35 | -9.1 |
| Participant 3 | 26.4 | -42.8 | -6.5 | -24.2 | -58.6 | -5.9 |
| Participant 5 | 23.1 | -58.6 | -5.9 | -27.4 | -58.6 | -5.9 |
| Participant 6 | 28.4 | -26.4 | -14 | -35 | -26.4 | -14.5 |
| Participant 7 | 26.4 | -47.3 | -5.9 | -22.1 | -47.3 | -5.9 |
| Participant 8 | 25.6 | -58.6 | -5.6 | -21 | -31.7 | -8.1 |
| Participant 9 | 21 | -31.7 | -11.3 | -26.4 | -31.7 | -10.2 |
| Participant 10 | 22.1 | -28.5 | -17.8 | -24.2 | -28.5 | -18.8 |
| Participant 11 | 25.3 | -42.5 | -8.1 | -32.8 | -42.5 | -8.1 |
| Participant 12 | 25.3 | -50 | -4.8 | -23.1 | -50 | -4.8 |
| Participant 13 | 23.1 | -35 | -8.1 | -26.4 | -35 | -10.2 |
| Participant 14 | 24.2 | -17.8 | -19.9 | -27.4 | -17.8 | -22.1 |
| Participant 15 | 25.3 | -50 | -5.9 | -21 | -50 | -7 |
| Participant 16 | 21.2 | -32.8 | -9.8 | -21.2 | -32.8 | -9.8 |
| Participant 17 | 23.1 | -31.7 | -9.1 | -22.1 | -31.7 | -9.1 |
| Participant 18 | 24 | -29 | -12.6 | -24 | -45.4 | -10.1 |
| Participant 19 | 24.2 | -43.6 | -7.2 | -25.3 | -42.5 | -7.2 |
| Participant 20 | 24.2 | -28.5 | -12.4 | -25.3 | -28.5 | -12.4 |
| Participant 21 | 26.4 | -41.4 | -8.1 | -24.2 | -40.3 | -8.1 |
| Participant 22 | 26.8 | -35.1 | -6.3 | -23.3 | -34.8 | -7.2 |
| Participant 23 | 28.5 | -37.1 | -7 | -25.3 | -38.2 | -7 |
| Participant 24 | 23.1 | -30.7 | -10.2 | -29.6 | -31.5 | -9.7 |
| **Average:** | **24.7** | **-37.9** | **-9.4** | **-25.3** | **-38.1** | **-9.6** |

### Behavioral Statistical Analysis

Performance and reaction time on the WM and recognition tasks were analyzed for 22 subjects in SPSS v24.0. Percent correct, d’ and reaction time (RT) were correlated with the group source waveforms in BESA Statistics v2.0 with corrections for multiple comparisons using non-parametric cluster-permutation testing (Maris & Oostenveld, 2007).

## Results

### Behavioral Results

There was a difference in accuracy (percent correct) between low-load (Mean = 90.64% correct, SD = 19.44) and high-load (78.27% correct, SD = 25.96) performance. A related-samples Wilcoxon Signed Rank test showed that there is a significant difference in performance accuracy between high- and low-load conditions (p=0.002), such that performance was better on the low-load task as compared with the high-load task. There was also a difference in reaction times between low-load (Mean = 855.40 msecs, SD = 197.13) and high-load (Mean = 882.50, SD = 279.82) conditions. A related-samples Wilcoxon Signed Rank test showed a significant difference in reaction times between high- and low-load conditions (p=0.033), such that reaction time was quicker on the low-load task as compared with the high-load task.

### Source Analysis Results

For group ERP source analysis weighted by fMRI data, low-vs-high load task types were compared for bilateral thalamic regions during the delay condition and bilateral parahippocampal gyrus regions during the encoding condition. *Figure 5* illustrates the time-varying cortical activity that explains the scalp ERP components and the slightly different dipole locations and orientations within the three-dimensional FEM head model of the delay and encoding conditions. The average Talairach coordinates are listed in *Tables 1 and 2* for both the loads under delay and encoding conditions, respectively.

**Figure 5:**
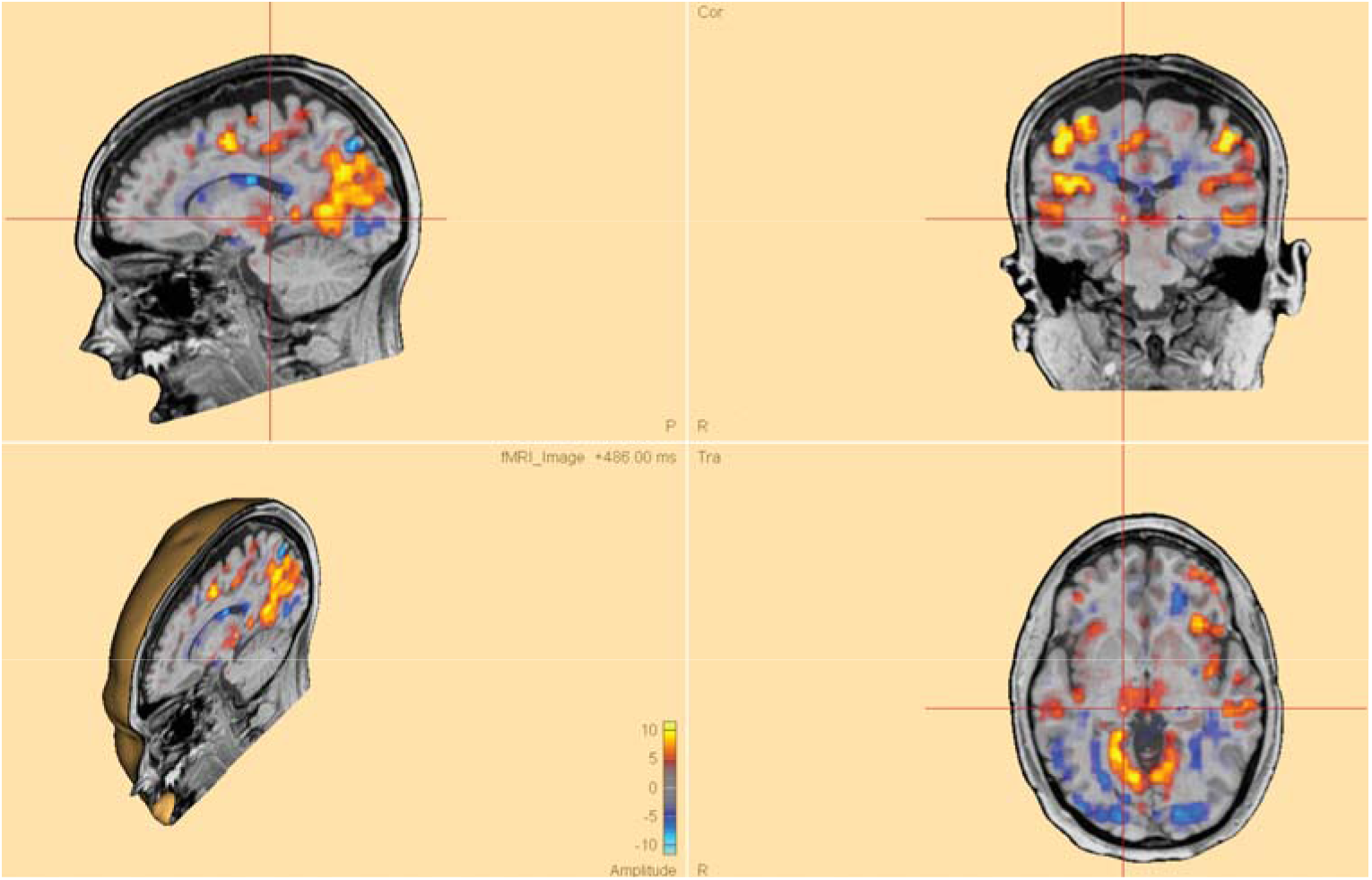
Dipole Position Example. Source Localization (*a priori* dipole modeling) highlighting the right (red; Talairach coordinates: x = −18.8, y = −25.3, z = 0.5) thalamus. Note that the realistic head model was created using the 4-layer Finite Element Model (FEM) as implemented in BESA v7.0. The warm colors (red to yellow) on the map reflect greater activation during delay period while the cold colors (blue) reflect greater activation during encoding period.

During the delay period, both left and right thalamus showed a WM load effect. Greater activation was observed in the bilateral thalamus during low-load delay when compared to high-load delay condition. The asymmetric dipole clusters for the left thalamus are shown in *Figure 6*. The source analysis results for high- and low-load conditions show a strong WM load effect where the activation is higher for low-load compared to the high-load delay condition between 160 msecs and 390 msecs (p=0.003).

**Figure 6.**
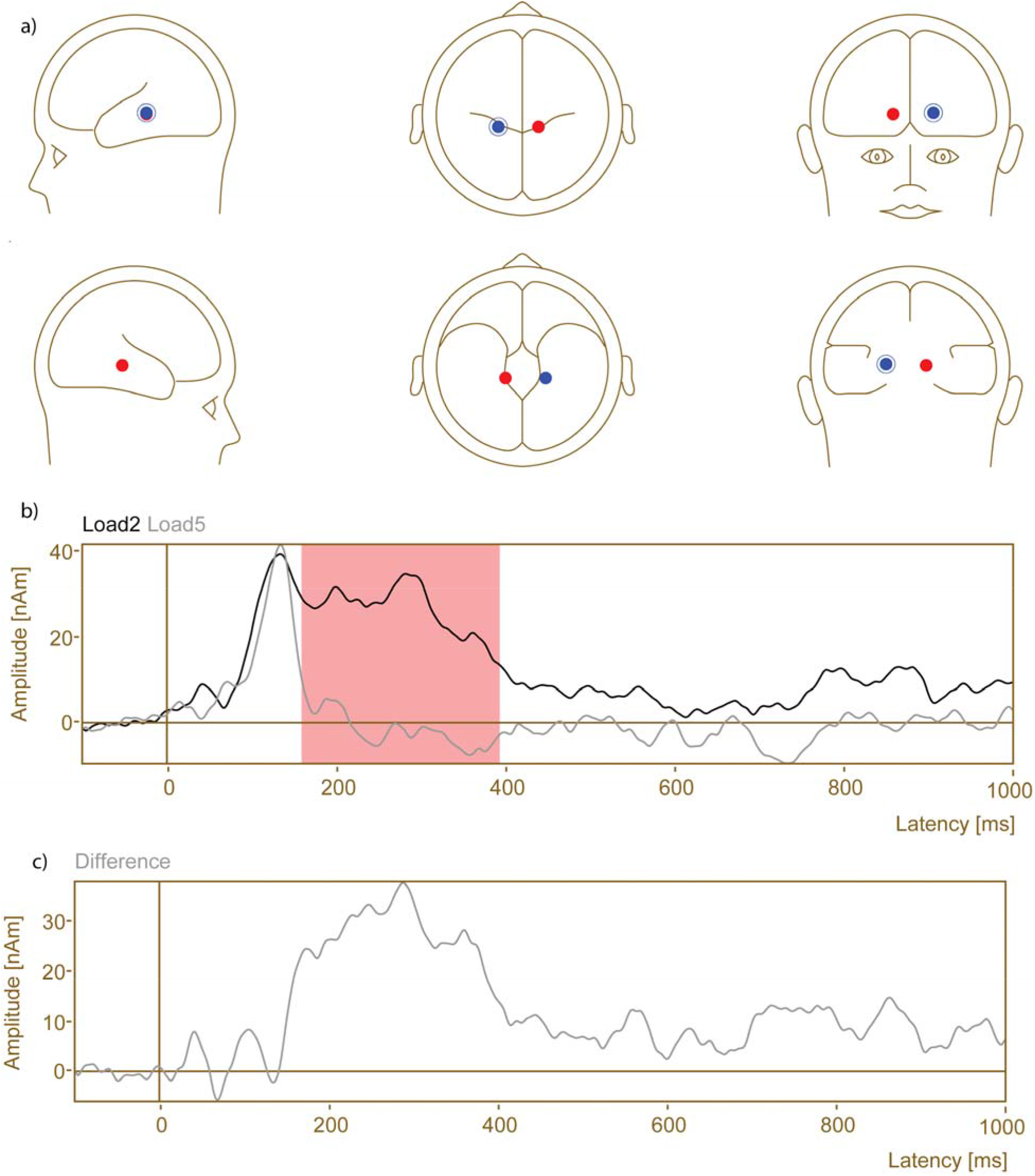
Group Source Waveform during Delay Activity Cluster 1. a) Head plot of asymmetric dipole clusters located in thalamus in left (blue) and right (red) hemispheres. [Cluster 1 (blue): p = 0.001; Cluster 2 (red): p = 0.024]. b) Source Analysis results for high (Load 5)- and low (Load 2)-load conditions for 1-dipole (left hemisphere) solution during delay (Baseline: - 100 msec, Delay Period: 0-1000 msec, collapsed across 6000 msec delay period). Group source-derived waveforms (grand average) of Load 2 (black) and Load 5 (gray) show a WM load effect occurring between 160 msec and 390 msec [p = 0.003]. c) Dipole source analysis delay period difference wave (Load 2 minus Load 5).

The asymmetric dipole clusters for the right thalamus are shown in *Figure 7*. The source analysis results for high- and low-load conditions show a statistically significant 4 clusters indicating WM load effect occurring in four different time intervals. The earliest significant effect is between 240 msecs and 330 msecs (p=0.022) followed by time intervals between 506 msecs and 582 msecs (p=0.034), 694 msecs and 760 msecs (p=0.046), and 858 msecs to 956 msecs (p=0.021). Similar to the delay comparison, activation is higher for low-load compared to the high-load condition.

**Figure 7.**
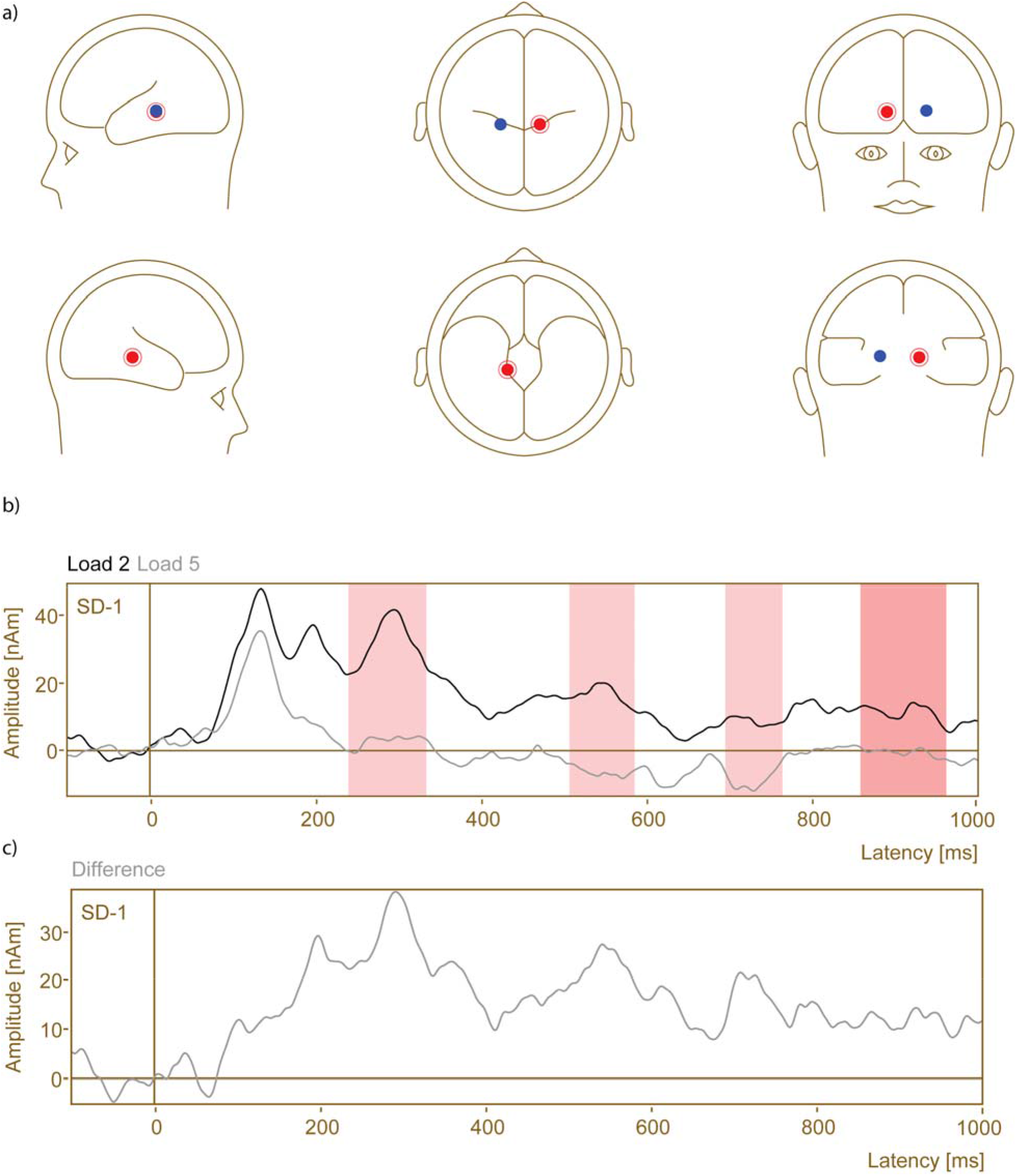
Group Source Waveform during Delay Activity Cluster 2. a) Head plot of asymmetric dipole clusters located in thalamus in left (blue) and right (red) hemispheres. [Cluster 2 (red): p = 0.021]. b) Source Analysis results for high (Load 5)- and low (Load 2)-load conditions for 1-dipole (right hemisphere) solution during delay. Group source-derived waveforms (grand average) of Load 2 (black) and Load 5 (gray) clearly show a WM load effect occurring between 240 msec and 330 msec, [p = 0.022], 506 msec and 582 msec [p = 0.034], 694 msec and 760 msec [p=0.046], and 858 msec to 956 msec [p = 0.021]. c) Dipole source analysis delay period difference wave (Load 2 minus Load 5).

For the encoding period, both left and right parahippocampal gyri did not show a significant WM load effect as the group source-derived waveforms did not show a significant difference between the two load conditions (p=0.486). The asymmetric dipole clusters for the left and right parahippocampal gyri are shown in *Figure 8*.

**Figure 8.**
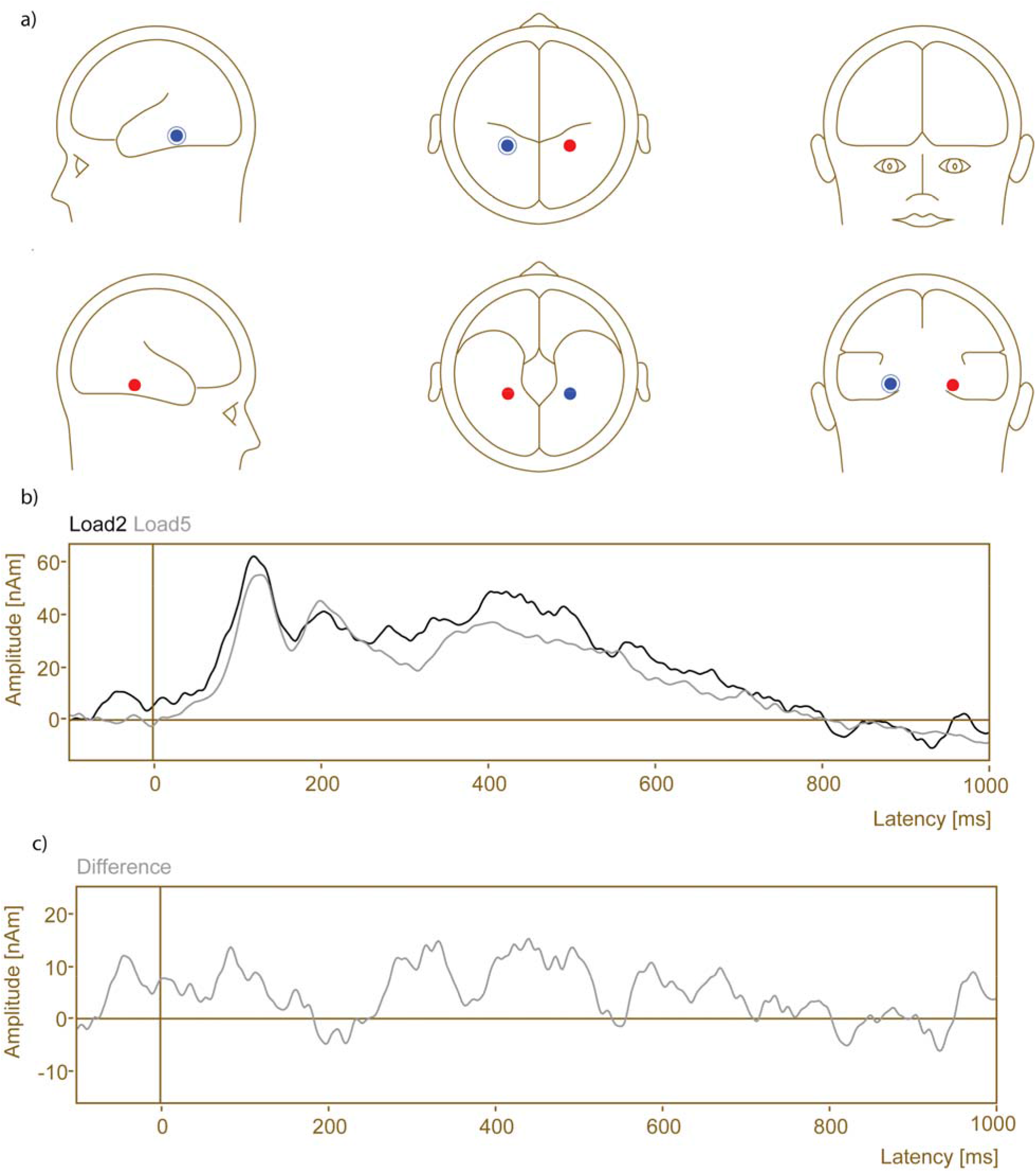
Group Source Waveform during Encoding. a) Head plot of asymmetric dipole clusters located in parahippocampal gyrus in left (blue) and right (red) hemispheres. [Cluster 1 (blue): p = 0.486]. b) Source Analysis results for high (Load 5)- and low (Load 2)-load conditions for 1-dipole (left hemisphere) solution during encoding (Baseline: - 100 msec, Encoding Period: 0 to 1000 msec of 1400 msec period). Group source-derived waveforms (grand average) of Load 2 (black) and Load 5 (gray) do not show a significant WM load effect [p = 0.486]. c) Results of the dipole source analysis of the difference wave during Encoding.

### fMRI Results

For fMRI analysis, group paired t-test difference maps were computed from BOLD fMRI data using 3dttest++ in AFNI for both delay and encoding conditions. In high vs. low load comparison during encoding period, right lingual gyrus extending anteriorly to parahippocampal gyrus, left middle occipital gyrus, left precentral gyrus, right thalamus, and left supramarginal gyrus regions show higher activation during high compared to low load (t > 3.67, p = 0.001, cluster size > 40 voxels). *Figure 9* shows fMRI activation differences as a function of load during encoding. *Table 3* lists the cluster sizes, coordinates, and brain regions with significant differences between high and low load during the encoding period.

**Figure 9.**
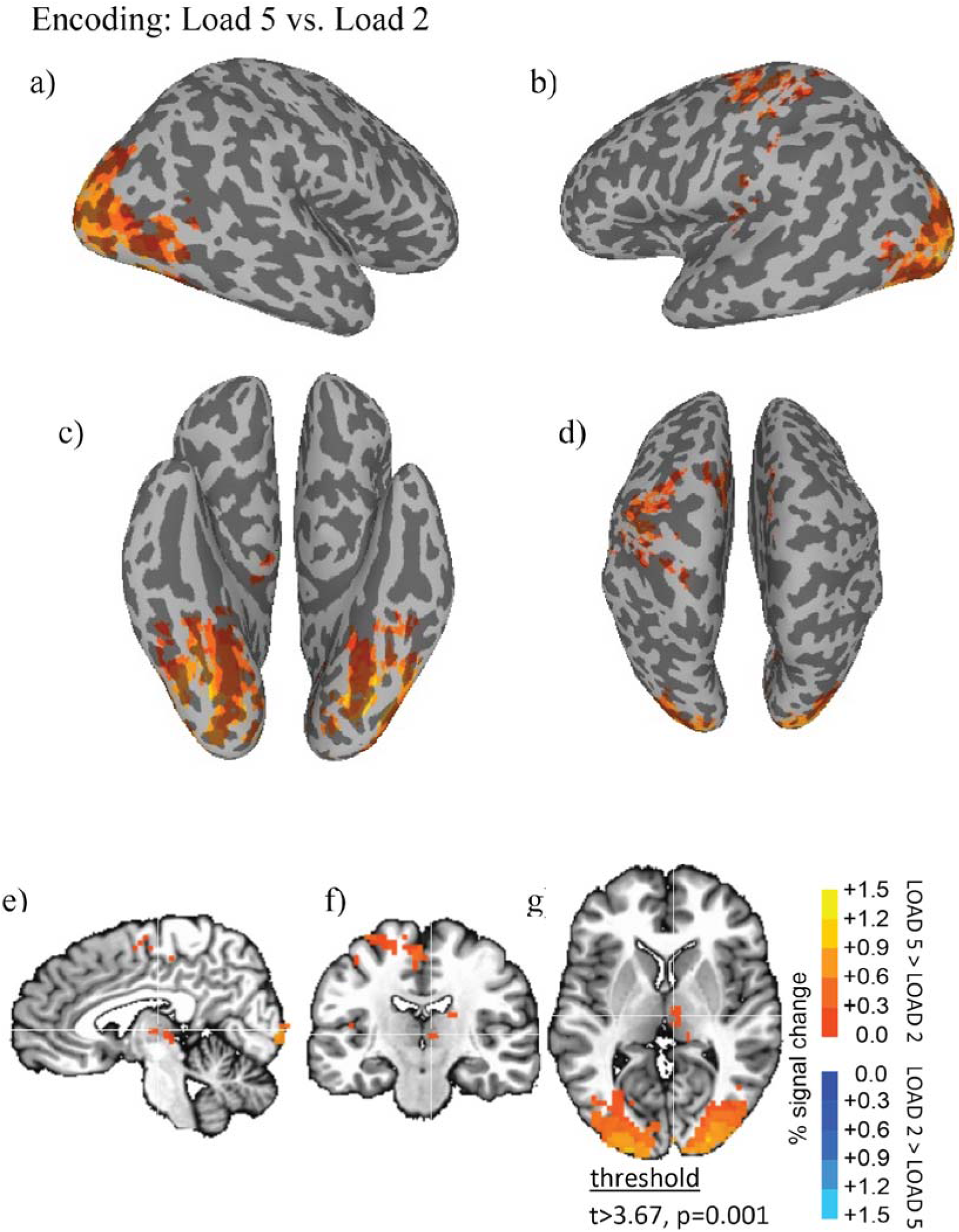
fMRI Results Show Brain Regions Differing Significantly Between High and Low Load During the Encoding Period. Thresholded fMRI statistical maps (t>3.67, p=0.001, cluster size > 40 voxels) displayed on inflated cortical surface representations (a = left hemisphere, b = right hemisphere, c = ventral view, d = dorsal view) and orthogonal views (e = sagittal, f = coronal, g = axial view). The crosshairs in the orthogonal view is located x=28, y=38, z=48 mm, the peak location of a cluster of activity in medial dorsal nucleus of the thalamus that was greater for high compared to low load encoding.

**Table 3.** Brain Regions with Significant Differences Between High and Low Load During the Encoding Period. Cluster sizes after thresholding at t>3.67 (p<.001) are reported as number of contiguous voxels in descending order. In parentheses after the cluster number it is indicated whether the activity in the cluster was greater at encoding during the high (L5) or low (L2) load condition. Here all five clusters showed greater activity during high load encoding compared with low load encoding. For each cluster, the x,y,z Talairach coordinate is reported in mm for the peak local maxima within the cluster followed by the labeled brain region.

| Cluster | Cluster Size | X | Y | Z | Brain Region |
| --- | --- | --- | --- | --- | --- |
| 1 (L5) | 1040 | 29 | -68 | -12 | R Lingual Gyrus |
| 2 (L5) | 921 | -31 | -89 | -6 | L Middle Occipital Gyrus |
| 3 (L5) | 297 | -31 | -14 | 63 | L Precentral Gyrus |
| 4 (L5) | 45 | 28 | 38 | 48 | R Thalamus |
| 5 (L5) | 45 | -46 | -23 | 18 | L Supramarginal Gyrus |

In high vs. low load comparison during delay period, right cuneus, left precentral gyrus, left supramarginal gyrus, right inferior frontal gyrus, right superior medial gyrus, right middle temporal gyrus, right middle cingulate cortex, right angular gyrus, and left thalamus regions show higher activation during low compared to high load (t > 3.67, p<.001). Additionally, left calcarine gyrus and left calcarine gyrus regions show greater activation during high compared to low load conditions during the delay period. *Figure 10* shows fMRI activation differences as a function of load during the delay period. *Table 4* lists the cluster sizes, coordinates and brain regions with significant differences between high and low load during the delay period.

**Figure 10.**
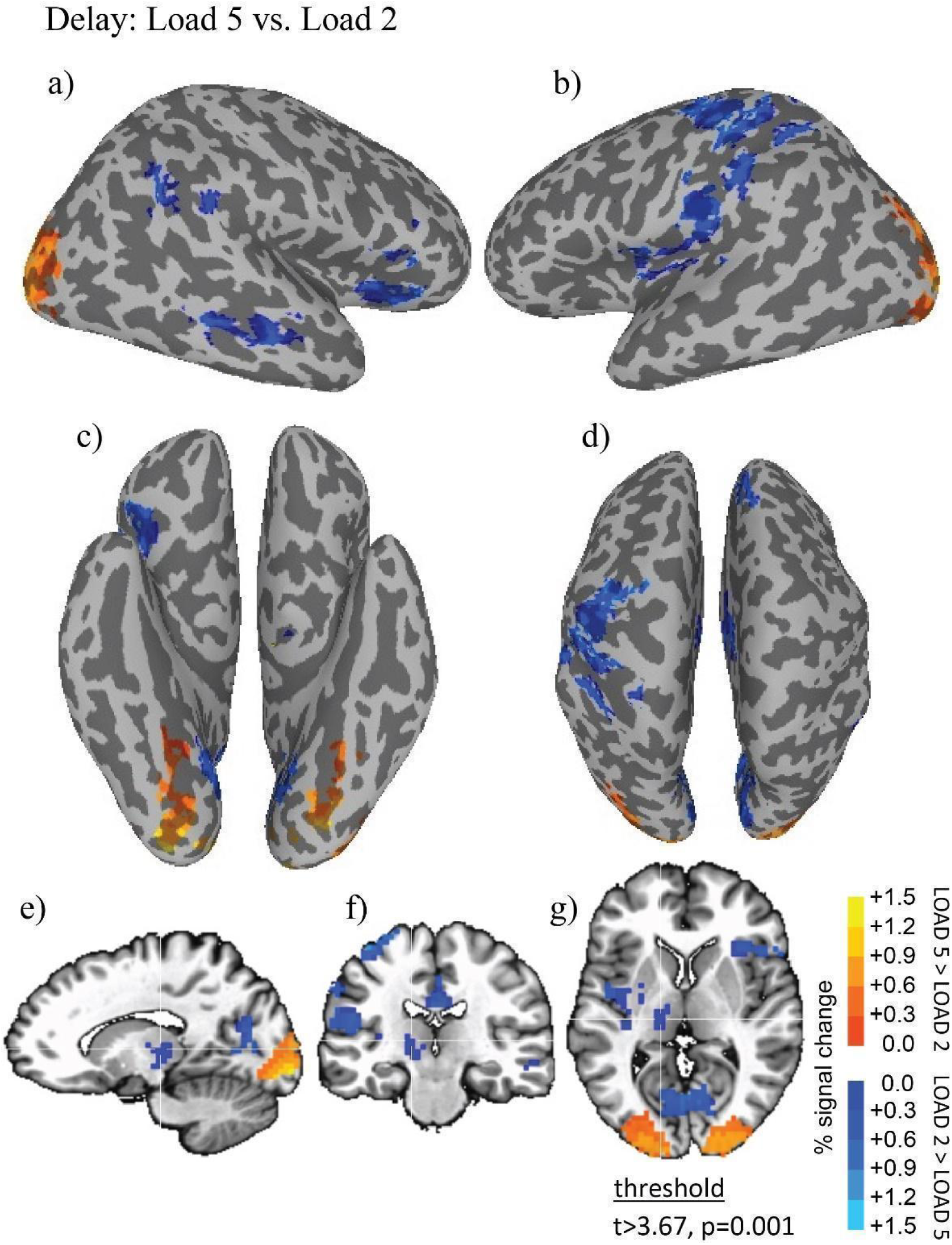
fMRI Results Show Brain Regions Differing Significantly Between High and Low Load During the Delay Period. Thresholded fMRI statistical maps (t>3.67, p=0.001, cluster size > 40 voxels) displayed on inflated cortical surface representations (a = left hemisphere, b = right hemisphere, c = ventral view, d = dorsal view) and orthogonal views (e = sagittal, f = coronal, g = axial view). The crosshairs in the orthogonal view is located x=-10, y=-17, z=12 mm, the peak location of a cluster of activity in medial dorsal nucleus of the thalamus that was greater for low compared to high load during the delay period.

**Table 4.** Brain Regions with Significant Differences Between High and Low Load During the Delay Period. Cluster sizes after thresholding at t>3.67 (p<.001) are reported as number of contiguous voxels in descending order. In parentheses after the cluster number it is indicated whether the activity in the cluster was greater in the delay period during the high (L5) or low (L2) load condition. In this comparison two clusters showed greater activity in high load delay period compared with the low load delay period, while 9 clusters showed greater activity in the low load delay period compared with the high load delay period. For each cluster, the x,y,z Talairach coordinate in mm is reported for the peak local maxima within the cluster followed by the labeled brain region.

| Cluster | Cluster Size | X | Y | Z | Brain Region |
| --- | --- | --- | --- | --- | --- |
| Cluster | Cluster Size | X |  |  |  |
| 1 (L5) | 472 | -10 | -92 | -6 | L Calcarine Gyrus |
| 2 (L2) | 447 | 2 | -68 | 9 | R Cuneus |
| 3 (L5) | 431 | 29 | -86 | -9 | R Inferior Occipital Gyrus |
| 4 (L2) | 421 | -37 | -26 | 63 | L Precentral Gyrus |
| 5 (L2) | 175 | -46 | -23 | 18 | L Supramarginal Gyrus |
| 6 (L2) | 127 | 44 | 19 | -3 | R Inf Front Gyrus (p. Orbitalis) |
| 7 (L2) | 120 | 5 | 31 | 57 | R Superior Medial Gyrus |
| 8 (L2) | 75 | 65 | -32 | -3 | R Middle Temporal Gyrus |
| 9 (L2) | 67 | 2 | -23 | 27 | R Middle Cingulate Cortex |
| 10 (L2) | 59 | 56 | -53 | 33 | R Angular Gyrus |
| 11 (L2) | 44 | -10 | -17 | 12 | L Thalamus |

### Correlation with behavior

As there were significant differences between loads during delay conditions in the bilateral thalamus, we tested whether the differential activation correlated with the behavioral performance. The individual subject grand source waveforms from both delay conditions were correlated with the percent correct performance. No statistically significant clusters that correlated with performance were found at either low (p = 0.475) or high load (p = 0.256). Individual subject grand source waveforms from both the delay conditions were also correlated with performance and reaction times, but no statistically significant clusters (p>0.05) were found.

## Discussion

The main finding in the present study was differential bilateral load-dependent thalamic activation during the delay period using simultaneous EEG-fMRI source analysis (*Figures 6 & 7*). There was no differential parahippocampal gyri activation during the encoding period as a function of memory load using simultaneous EEG-fMRI source analysis (*Figure 8*). On the contrary, only fMRI analysis showed a significant difference during encoding period (*Figure 9*). During encoding, fMRI results showed higher activity in left middle occipital gyrus, left precentral gyrus, left supramarginal gyrus, right thalamus, and right lingual gyrus, during high load when compared to low load conditions. Load related differences during encoding were centered on lingual gyrus, which is located more posterior than parahippocampal gyrus, slightly contrary to our initial hypothesis. During the delay period, fMRI results showed higher activity in right cuneus, left precentral gyrus, left supramarginal gyrus, right inferior frontal gyrus, right superior medial gyrus, right middle temporal gyrus, right middle cingulate cortex, right angular gyrus and left thalamus during low compared to high load conditions. Additionally, fMRI analysis revealed higher activation in left calcarine gyrus and right inferior occipital gyrus during high load when compared to low load conditions. It noteworthy that fMRI analysis revealed thalamic activation that was lateralized to the left hemisphere during the delay period. fMRI analysis also revealed higher visual cortex activation during the high-load delay period. The behavioral data (rate of correct responses and reaction times) were used to investigate if there were any correlation with the source waveforms during the delay periods and performance. However, no significant correlations were found between thalamic activation and performance. This lack of significant correlation is perhaps because of overall differences in behavioral performance between the low and high load conditions.

Previous studies have linked WM delay activity in the posterior and prefrontal brain regions associated with a limited capacity WM buffer, where maintaining more items in the buffer leads to an increase in activity (Rypma et al., 1999; Schon, Quiroz, Hasselmo, & Stern, 2009). The prefrontal cortex (PFC) has been implicated in WM delay activity, suggesting that it is involved in item maintenance during delay activity (Inagaki et al., 2019). The PFC receives connections from the thalamus, which has been shown to play a critical role in working memory (Guo et al., 2017). Few studies have looked at load effects in the thalamic areas during delay activity. In our current research, WM load effects occurred in bilateral thalamus during delay activity using simultaneous EEG-fMRI source analysis. The differential activation in the bilateral thalamus was such that lower activation corresponded to high-load, while greater activation corresponded to low-load condition. On the other hand, the corresponding fMRI results showed higher activation in the visual cortex during high-load delay periods when compared to low-load conditions. The finding suggests that the brain could be relying more on input from subcortical regions like the thalamus when less information is being maintained in WM (e.g., during low load) and could be relying more on sensory areas like the visual cortex when more information is being maintained in WM (e.g., high load).

The sensory gating mechanism could explain the differential activation of the thalamus. The thalamus could be regulating the gain of sensory processing during the different load conditions such that the processing of potentially disrupting sensory information (scrambled images) is down-regulated when this information might be interfering with the retained information as in high-load condition. On the other hand, sensory processing of scrambled images is upregulated when this information might not be potentially interfering with the retained information during the low-load condition. The detection of the interfering stimuli could be triggering activity in the dlPFC via thalamic relays (Knight et al., 1999). PFC has been conceptualized as a dynamic filtering mechanism as well (Shimamura, 2000). The thalamus could be applying a similar dynamic filter to retrieve and select information relevant to the current task requirements. The reciprocal connections between the thalamus and PFC could be influencing such activity in concert. There is evidence that dlPFC is involved in goal-based control by inhibition of task-irrelevant information (Ridderinkhof, Van Den Wildenberg, Segalowitz, & Carter, 2004). Using transcranial magnetic stimulation (TMS), goal-based representations in PFC were used to modulate how perceptual information is selectively filtered such that the task goal, specified by the instruction, can modulate perceptual processing by inhibiting task-irrelevant information (Feredoes, Heinen, Weiskopf, Ruff, & Driver, 2011). Thalamic projections to the PFC are suggested to inhibit task-irrelevant information in the context of cognitive control.

In demanding N-back working memory tasks, the dlPFC network expands, showing marked connectivity with parietal regions and areas in the ventral visual pathway (Cohen & D’Esposito, 2016). With greater working memory demands, the thalamus may signal the PFC to increase the connection strength of item representations to become greater within networks, including the parahippocampal regions, parietal regions, and areas in the visual cortex.

We hypothesize that the thalamus may be enhancing the task-relevant information or inhibiting task-irrelevant information. On this account, during the high-load condition, the thalamus may serve to inhibit distracting information to activate relevant stimuli information in higher cortical areas like the primary visual cortex. This account would help explain our results from the study combining both fMRI and EEG methods. The thalamus may be involved in successfully orchestrating inhibitory control when high-load information is being maintained. Therefore, the thalamic responses could be attenuated to suppress the complex environmental stimuli. The differential thalamic activation could also explain the difference in behavioral performance in both loads. The highest evoked activity during maintenance of fewer stimuli in the presence of interfering perceptual stimuli implies better consolidation of stimuli, which eventually leads to better performance (Lavie, Hirst, De Fockert, & Viding, 2004). Alternatively, the thalamus, with its reciprocal connections with PFC and motor areas, may serve to prepare the participant for the appropriate behavioral response (Fonken, 2016). On this account, higher thalamic activation could mean more increased preparedness and readiness to make a behavioral response during the low-load condition. This account would also explain that lower thalamic activation would imply reduce confidence and readiness to make the relevant behavioral response because more scenes had to be maintained during the higher load condition.

It is to be noted that we originally did not intend to study interference per se during the delay period. The scrambled images presented during the delay were chosen to serve as a perceptual baseline with similar color and spatial frequency for comparing brain activations between encoding and delay conditions. However, the scrambled images serve as interfering stimuli as subjects must maintain the previously presented intact scenes in the face of scrambled visual input during the delay. This presumably becomes more difficult as more stimuli presented during encoding must be maintained in the face of scrambled input during the delay. It is noteworthy and surprising that greater thalamic source activity was associated with lower load, which is counterintuitive to many hypotheses that usually associate higher activity with higher load.

### Limitations of the Current Study

Potential limitations of the present study include some caveats when comparing significant statistical results between EEG and fMRI methodologies. The differences in EEG and fMRI results could be explained by the use of separate statistical software packages for analyses, with EEG analysis relying on non-parametric permutation testing to determine threshold and fMRI analysis relying on more traditional parametric methods in AFNI. For our EEG data analysis, we used BESA Statistics, which uses a non-parametric cluster-permutation method (Maris & Oostenveld, 2007). For our fMRI analysis, we used AFNI, which uses a parametric approach (Cox, Chen, Glen, Reynolds, & Taylor, 2017). The stronger fMRI activations in the high load condition could be because more data points (more stimuli presented during high versus low load encoding) went into the analysis, which means more data and time input for fMRI signal averaging.

Another potential limitation is slight inaccuracies for the EEG electrode position based on coregistration with MRI scans. We initialized positions using a standard electrode montage instead of individual digitized electrode positions due to lack of a digitizer tool at the MRI facility. While imperfect electrode localization estimates could have compromised the quality of the source analysis, individual adjustment of electrode positions were made based on visible electrode indentations on the scalp surface reconstructions. Additionally, the number of channels used for the study was 32 instead of 64 or 128 due to subject participation and set-up time limitations. The limited number of channels could have resulted in a less precise source analysis result (Song et al., 2015). However, compared with the scalp recorded ERPs, our source analysis results closely matched our fMRI results. Scalp recorded ERP topographies for the delay and encoding conditions are shown in *Supplemental Figures 1 and 2*, respectively.

### Future Studies

Future studies could examine how differential thalamic activation and connectivity relates to behavioral performance under a range of different WM loads to equate performance. Further research will also be necessary to determine the degree to which WM maintenance mechanisms reflect the selection of task-relevant information versus the inhibition of irrelevant information. This idea could be tested using category-specific stimuli, where the participants would be asked to remember or ignore specific categories during each trial. Furthermore, it would also be interesting to test for differences in thalamic activations as a function of delay period length. By increasing the delay period duration, one could find evidence whether the thalamic activation rises just before the behavioral response when the end of the delay period is unpredictable. Future studies should further test whether there is truly a negative correlation between thalamus during the delay activity and posterior cortical areas during encoding and its relation to successful retrieval of items.

## Conclusions

The study results support the view that the thalamus contributes to complex cognitive functions like memory maintenance as opposed to the traditional view of the thalamus as serving as a simple sensory relay system. The results suggest that thalamus is involved in working memory and is differentially active as a function of WM load. When the WM load is low, the thalamus is more active than when WM load is high. During high load, the thalamus may function to attenuate incoming distracting perceptual stimuli while during low load the thalamus shows less attenuation in the face of distracting perceptual input. This suggests that the thalamus is preparing the superficial cortical areas for successful task-relevant information during high load WM maintenance. More research is needed to understand how the thalamus and PFC work in concert to inhibit potentially disruptive or irrelevant information and modulate attention during maintenance.

## Acknowledgements

We thank Ahmed Duke Shereen, Ph.D. and Kenneth Ng for help task set-up and data collection. We thank Farzana Antara, Gabriela Silva, Albert Perdomo, and Aliza Hacking for help with EEG preparation and analysis.

## Declarations

### Availability of data and material

Deidentified data will be made available upon reasonable request to the corresponding author.

### Code availability

No code was developed to perform these analyses. All analyses were performed using widely available and validated data analysis packages as described in the methods section.

## Supplementary Information

**Supplemental Figure 1:**
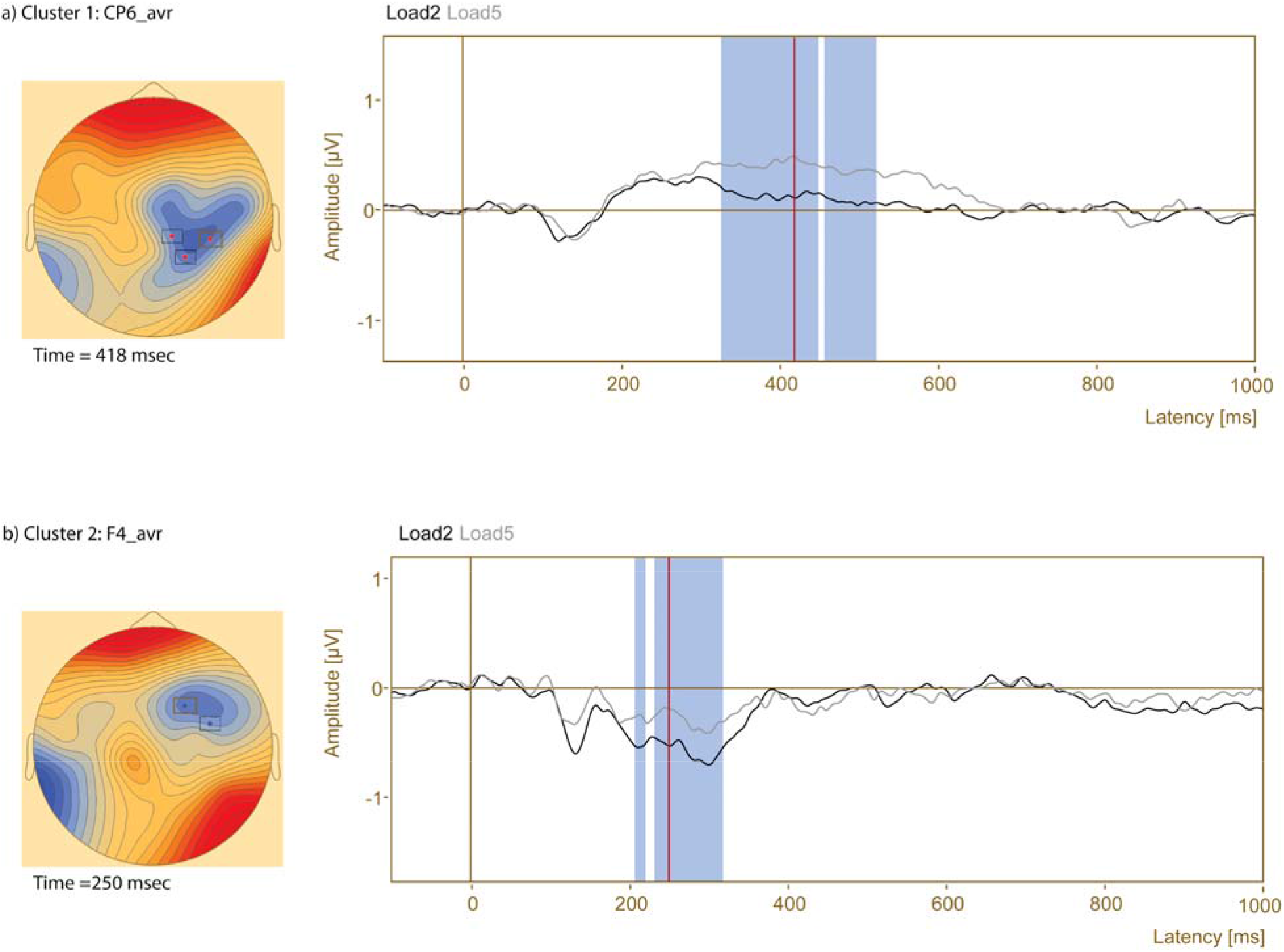
High vs. Low Load Grand-Average Sensor-Level ERP Waveform and Topography Comparison during the Delay Period. Sensor level analysis was done on 22 subjects to show the scalp topography during high- and low-load delay conditions. Significant differences occurred at a) right centro-parietal sensor location between approximately 326 msecs and 518 msecs (p=0.008), where Load 5 (gray) amplitude is greater than Load 2 (black) amplitude; a b) right frontal sensor location between approximately 198 msecs and 344 msecs (p=0.023), where Load 2 (black) amplitude is more negative than Load 5 (gray) amplitude. The top down views (left) correspond to the most significant time-point, indicated by the red solid lines. The highlighted blue shaded regions indicate more negative amplitude for load 2 than Load 5.

**Supplemental Figure 2:**
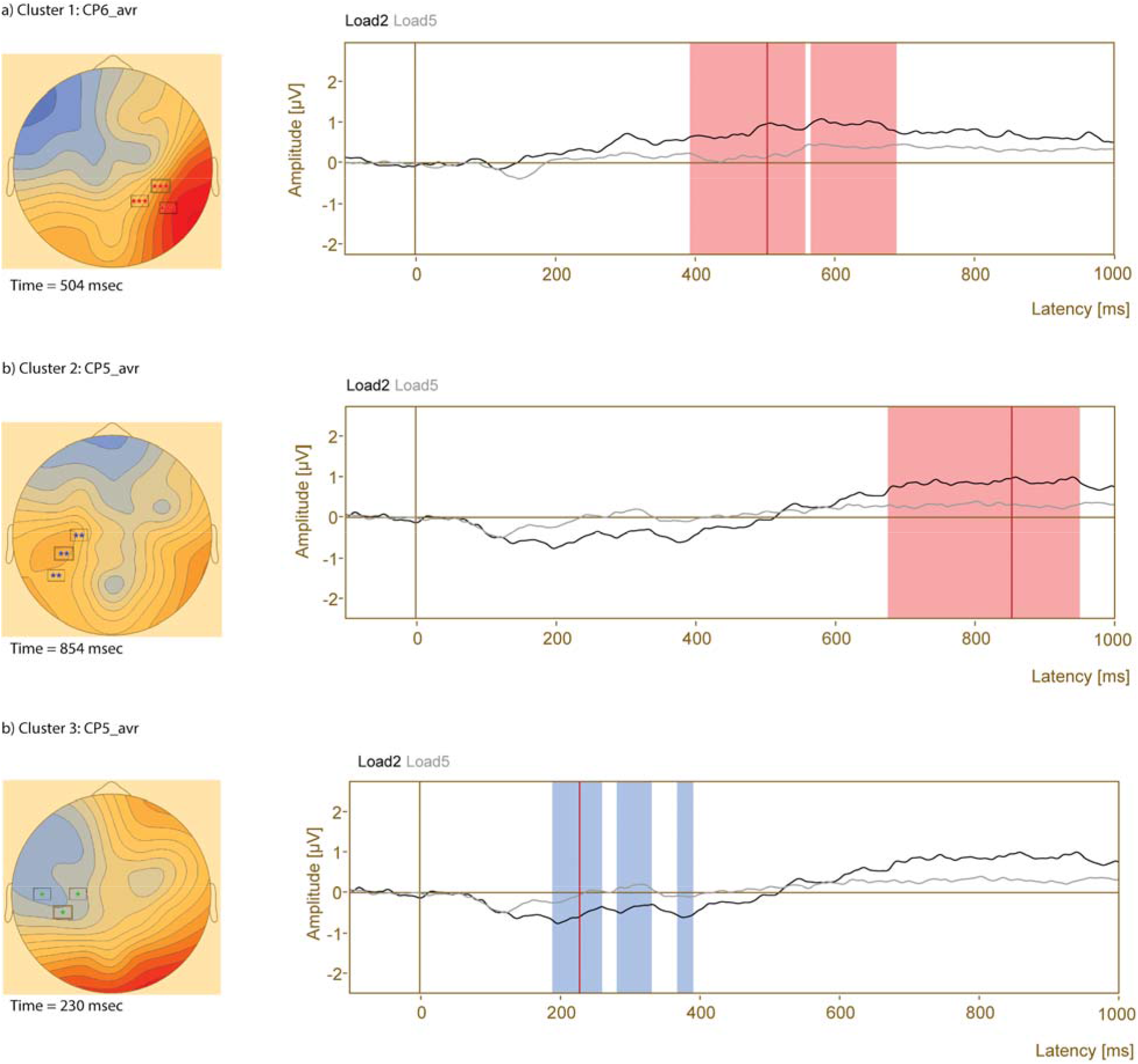
High vs. Low Load Grand-Average ERP Sensor-Level Waveform and Topography Comparison during the Encoding Period. Sensor level analysis was done on 22 subjects to show the scalp topography during high- and low-load encoding conditions. Significant differences occurred at a) right centro-parietal sensor location between approximately 252 msecs and 684 msecs (p=0), where Load 2 (gray) amplitude is greater than Load 5 (black); b) left centro-parietal sensor location between approximately 618 msecs and 962 msecs (p=0.002), where Load 2 (gray) activation is greater than Load 5 (black); c) left centro-parietal sensor location between approximately 190 msecs and 476 msecs (p=0.007), where Load 2 (gray) activation is greater than Load 5 (black). The top-down views (left) correspond to the most significant time-point, indicated by the solid red lines. The highlighted blue shaded regions indicate more negative amplitude for Load 2 than Load 5, while the red shaded regions indicate more positive amplitude for Load 2 than Load 5.

